# Towards MR-based interrogation of the hypoxia-driven insulin resistance mechanism: Adipocytes size estimation

**DOI:** 10.1101/2025.05.29.656865

**Authors:** Darya Morozov, Isaac Prentiss, Nelson Yamada, Sasha Hakhu, Alexander L. Sukstanskii, Scott C. Beeman

## Abstract

Obesity is a major risk factor for type 2 diabetes, yet not all individuals with obesity develop metabolic disease, underscoring the need for mechanistic biomarkers. Adipocyte hypertrophy is a hypothesized driver of insulin resistance, but current methods for quantifying adipocyte size are invasive. Here, we propose and validate a non-invasive MRI approach based on “short diffusion time” diffusion-weighted MR spectroscopy to estimate adipocyte size in vivo. Monte Carlo simulations confirmed the method’s accuracy across a physiologic range of adipocyte sizes (20 - 150 μm) and signal-to-noise ratios (SNR > 40). We applied this technique to the epididymal white adipose tissue (eWAT) of rats using in vivo 4.7T and ex vivo 11.7T MRI. Adipocyte sizes derived from diffusion MRI showed good agreement with histology, with minor systematic underestimation corrected by empirical factors. This approach does not require complex modeling or high diffusion weighting, increasing its translatability to the laboratory and clinical settings. Diffusion MRI may serve as a non-invasive “virtual biopsy” to monitor adipocyte morphology and improve understanding of obesity-related metabolic dysfunction.

## 1. Introduction

Obesity is a growing worldwide epidemic [1–4]. While it is clear that obesity is associated with a number of serious downstream complications, for example type 2 diabetes (T2D) it is not always clear by which mechanism(s) obesity drives these downstream pathologies nor is it entirely clear that obesity alone translates to unfavorable outcomes [5]. Indeed, this point is punctuated by the fact that roughly 25% of people with obesity are otherwise metabolically normal as determined by key clinical metrics like insulin sensitivity, blood pressure, liver fat, blood triglycerides, and hemoglobin A1c (HbA1c) [6, 7]. The existence of this dichotomy suggests that it is not obesity, per se, that causes diseases like T2D, but instead the complex microstructural and biochemical changes that obesity can induce in some people but not others. One such obesity-related microstructural change that can influence metabolic health is white adipocyte size [8, 9].

White adipocytes, the body’s primary lipid and energy storage depots, are very dynamic cells that are able to shrink and grow based on caloric intake and systemic energy expenditure. The typical diameter of an adipocyte in a lean, metabolically normal person is roughly 60 μm [10] whereas in case of obesity adipocyte diameters can exceed 150 μm [9]. Adipocyte hypertrophy of this magnitude can have the dual effect of decreasing the effective capillary density of the adipose tissue and increasing the transit distance required for oxygen to diffuse from the blood to the mitochondria, thus inducing a state of hypoxia [9]. Adipose hypoxia has been shown to trigger a shift from aerobic metabolism to anaerobic glycolysis [11, 12], adipose inflammation/fibrosis [13, 14], and ultimately insulin resistance [15]. Thus, adipocyte hypertrophy is hypothesized to be a primary initiator of insulin resistance [16–19]. This hypothesis has been investigated via postmortem- and biopsy-based and histologic methods, though methods for measuring adipocyte size in vivo are lacking. A non-invasive method for quantifying adipocyte size in vivo would allow for dynamic monitoring of adipocyte size in parallel with transient clinical metrics like insulin sensitivity, blood pressure, HbA1c, and liver fat. Such a method would be a major advance towards resolving causal mechanisms of the pathogenesis of type 2 diabetes. We hypothesize that magnetic resonance imaging (MRI) may provide a means to these ends.

MRI has proven itself as an efficient, non-invasive, quantitative and longitudinal tool for characterization of tissue biophysical and functional properties both in vivo and ex vivo. Traditional MR techniques to study adipose tissue use chemical shift Dixon-based imaging methods to assess the fat fraction in the tissue [20–25]. More recently, attempts towards characterization of lipid droplet size distribution by diffusion MR have been made by modeling the restricted diffusion behavior of lipids within a droplet [26–28]. These methods are based on Murday and Cotts approach describing restricted diffusion signal behavior within spherical boundaries [29]. In the framework of this approach, the signal is presented in the form of the infinite series, each term obtained from the corresponding term in the eigenfunction expansion of the well-known diffusion propagator within a sphere with impermeable boundaries [30]. Recently, this method was implemented to study small lipid droplets like those usually found in brown adipose tissue or in skeletal muscle [27, 31] and to measure large lipid droplet sizes found in tibia bone marrow in vivo using diffusion-weighted MRS at 3T [28]. The shortcoming of this technique is a slow convergence of the series at short diffusion times. In this regime, an average displacement of diffusing particles is much smaller than the actual droplet’s size and, as result, the area within the boundaries is not fully probed. Therefore, to accurately estimate the lipid droplet size using the Murday and Cotts approach, one is required to use a large number of terms included in the aforementioned eigenfunction series.

The alternative approach that can be used to estimate the lipid droplet size is the so-called “short time regime” diffusion MR. This regime takes place when an average displacement of the diffusing particles over the observation time is significantly smaller than the actual compartment size [32, 33] (see “Theory” section below). The advantage of this method over the Murday and Cotts approach takes the form of a simple diffusion-weighed MR signal model that does not require the eigenfunction expansion to estimate the size of the sphere. An additional advantage of this approach is that “short time regime” diffusion MR requires relatively low diffusion weighting, which is important for its clinical applications.

Herein, we hypothesize that fast, direct, and clinically accessible in vivo measures of adipocyte size are achievable using “short time regime” diffusion MR. We first examine the approach using Monte Carlo simulations and then apply it to in vivo and ex vivo studies of epididymal white adipose tissue of rats.

## 2. Materials and Methods

### 2.1 Theory

The vast majority on diffusion MRI experiments are performed in the long diffusion time regime. In this regime, an average displacement of spin-bearing molecules (usually, water) is much larger than a compartment’s size, and the characteristic diffusion time t_D_=R^2^/6D_0_ (*R* is the system’s size, *D*_0_ is the molecules’ diffusivity, and the factor of 6 accounts for diffusion in three dimnesions) is much shorter than the observation time. Indeed a typical cell size might be on the order of 10 – 30 microns and the diffusivity of water is ∼2 um^2^/ms, thus the time that it takes for the average water-bound proton to sample the diameter or a cell is in the ∼8 - 75 ms range, whereas a typical obseration time is on the order of 50-100 miliseconds (hardware limitations). As a result, over the course of a single MR observation, diffusing molecules will have sampled the entire cell volume, encountering and “bouncing off” of the boundaries of the compartment (cell).

The situation is quite different in the case of triglyceride molecules (*D*_0_ ∼ 0.02 µm^2^/ msec, see below) and the relatively large size of white adipocyte droplets (∼30 to 150 µm) in human adipose tissue. Even for the smallest droplets, the characteristic diffusion time is on the order of 7 sec or more, a timescale that is much longer than MRI measurement time. Thus, lipid diffusion within an oil droplet is an ideal application of short-time regime theory for quantifying compartment sizes; i.e., oil droplet size and thus white adipocyte size, which are typically comprised 99% of a single droplet.

In the short time regime, the molecules within a system are effectively partitioned into two pseudo-compartments. In one pseudo-compartment, which is proximal to the system’s boundaries, diffusion is restricted, whereas in the second pseudo-compartment, which is distal to the boundaries, diffusion is effectively free (molecules do not encounter the boundaries). The fractional populations of these pseudo-compartments propagate into the diffusion-attenuated MRI signal as fractional signal amplitudes which can be extracted either directly via very high b-value MRI data and a biexponential signal model (a method which demands extremely high MRI pulsed field gradient performance) or indirectly, using low b-value data and a mono-exponential signal model. It is the latter approach that will be explored in this work.

For the low *b*-value case explored herein (*bD*_O_ << 1), the diffusion-attenuated MR signal can be well described by a mono-exponential function [34]:

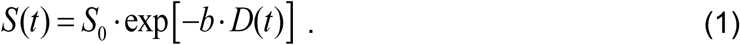

Here *S*_0_ is the signal amplitude, *b* is the *b*-value and *D*(*t*) is the time-dependent apparent diffusion coefficient (ADC). For a Stejskal-Tanner pulse sequence with the parameters Δ and δ the *b*-value is determined by:

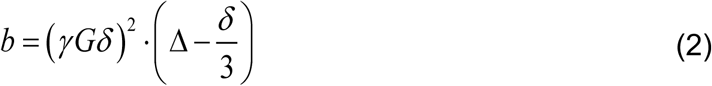

where γ is a gyromagnetic ratio, *G* and δ are the amplitude and the duration of the diffusion-sensitizing gradient, repectively, and Δ is the diffusion time.

It should be noted that Eq. (1) does not take into account the longitudinal relaxation. It is justified when performing *in silico* analysis (Monte Carlo simulations, see below). However, in the “real world” scenarios of *in vivo* and *ex vivo* diffusion-weighted Stimulated Echo Acquisition Mode (STEAM) MRI (see below), longitudinal relaxation must be accounted for. This may briefly be explained as an artifact of the STEAM sequence in which the magnetization is tilted to the z-direction for storage during the time over which proton diffusion is encoded, Δ. During this time, all protons are subject to longitudinal relaxation. When estimating *D*(*t*) from data with multiple *b*-values and a single diffusion time Δ, longitudinal relaxation can be ignored because the diffusion time (and thus time for longitudinal relaxation) is the same for all *b*-values. In our *in vivo* and *ex vivo* experiments, we use data with multiple *b*-values and multiple diffusion times, thus the longitudinal relaxation rate constant of white adipose lipid (*R*_1_) can be accounted for as follows:

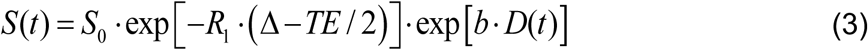

where *TE* is the input echo time (this quantity is the same for all experiments regardless of diffusion time Δ).

At sufficiently short diffusion times (*t* << *t_D_*), *D*(*t*) can be related to the surface-to-volume ratio (*S* / *V*) of the system [33, 35–37]:

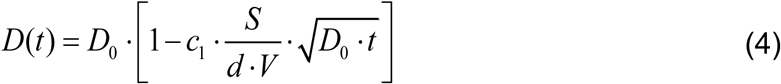

where *d* is the system’s dimensionality (*d*=3 for diffusion within a sphere), and the coefficient *c*_1_ depends on a gradient pulse sequence. For a Stejskal-Tanner pulse sequence with the parameters Δ and δ, *t* = Δ + δ, the coefficient *c*_1_ is equal to [36]:

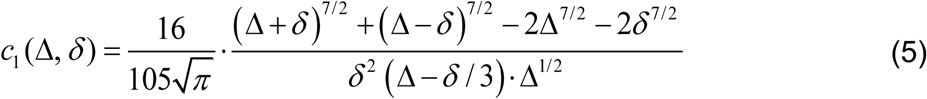

In the short gradient pulse approximation, δ << Δ, the coefficient *c*_1_(Δ, δ) reduces to Mitra’s value of 4/ 3√π [33].

In our case, when the lipid droplet within adipocyte cell is represented as a sphere with size *L*,

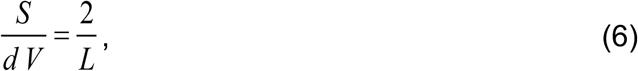

and Eq. (4) can be rewritten as

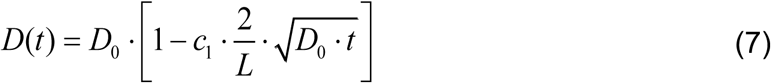

### 2.2 Simulations

To check the accuracy of the “short-time-regime” method for determining the size of compartments filled with slowly diffusing particles, Monte Carlo (MC) simulations were performed. The parameters of MC simulations (10^6^ diffusing particles with random starting positions) are as follows: the free diffusion coefficient *D* = 0.0175 µm^2^ /ms, diffusion time step 0.001 ms, gradient duration δ = 4 msec, diffusion time Δ ranges from 50 to 400 ms, and spheres’ size (*L*) are 20, 30, 40, 50, 60, 80, 100, 120 and 150 μm.

Gaussian-distributed noise was added to simulated (noiseless) diffusion signal decay in order to determine the effect of limited signal-to-noise ratio (SNR) on estimation of parameters The SNR values were 40, 60, 80, 100 and 200.

### 2.3 MR experiments

#### 2.3.1 In-vivo eWAT

All animal procedures were approved by the Washington University Division of Comparative Medicine. Three male Sprague Dawley Rats (n=3, Charles River Lab, USA) having weights of ∼400 grams were scanned.

All MR experiments were conducted at 4.7T MRI (Agilent Technologies, Santa Clara, California, USA) using a lab-built actively decoupled, transmit/receive volume coil. Rats were anesthetized with 2% isoflurane and placed on their back into the volume coil. Respiration was monitored throughout the experiment to ensure that animals maintained a ∼45 breaths per minute respiration rate. Animal temperature was kept constant on 37°C by blowing temperature-controlled air over the animal while monitoring core temperature with a rectal thermometer.

Diffusion experiments were respiratory-triggered and collected using a custom diffusion-weighted localized Stimulated Echo Acquisition Mode (STEAM). A 5 x 5 x 5 mm^3^ isotropic voxel was placed inside visceral (epidydimal) fat and short-time regime diffusion data were acquired with following parameters: repetition time (TR)/ echo time (TE)=2500/22 ms dummy scans (DS) = 4, number of averages (NA) = 1, number of repetitions (NR) = 8. The following diffusion parameters were used: Δ =50, 100, 150, 200 and 400 ms, δ = 4 ms, *b*-value range from 1.5 to 20 ms/µm^2^ (8 *b*-value steps), signal-to-noise ratio (SNR) ∼ 150 at *b* = 1.5 ms/µm^2^ and Δ = 50 ms. To cancel unwanted cross terms, data were acquired with diffusion-sensitizing gradients of both positive and negative polarities in three orthogonal directions (±x, ±y and ±z). Next, the geometric mean between positive and negative sets of experiments in the same direction for each targeted *b*-value was computed and averaged between three orthogonal diffusion directions and used in further diffusion analysis.

#### 2.3.3 Ex-vivo eWAT

After in-vivo scan, the animals were sacrificed and the epidydimal fat pads were resected and fixed in 10% PFA buffer solution for at least 24h prior to MR scan. Formalin-fixed rat eWAT tissues (n=3) were placed in a 12-mm glass tube filled with Fluorinert parallel to the B_0_ (z direction). All samples were temperature equilibrated to 37°C for a minimum of one hour prior to MR experiment.

All MR experiments were conducted at 11.7T (Agilent Technologies, Santa Clara, California, USA) using custom-build transmit/receive quadrature coil and a custom diffusion-weighted STEAM sequence. The of 3 x 3 x 3 mm^3^ isotropic voxel inside the tissue was selected the short diffusion time regime data were collected on using the following parameters: TR/TE=1500/17 ms, DS of 16, NA of 1, NR of 5. The short-diffusion-time regime was implemented with following experimental parameters: Δ = 50, 100, 150, 200 and 400 ms, δ = 4 ms, *b*-value range from 1.5 to 20 ms/µm^2^ (10 b-value steps), *SNR* ∼ 2500 at *b* = 1.5 ms/µm^2^ and Δ = 50 ms.

### 2.4 Data analysis

#### 2.4.1 Image and spectra post-processing

**In-vivo and ex-vivo eWAT tissues**: The diffusion signal amplitudes of the methylene-associated peak (1.3 ppm) were estimated for each repetition using the time domain data analysis in Bayesian probability theory-based software developed in our laboratory. (A toolbox of Bayesian-based, data analysis programs is available for free download at http://bayesiananalysis.wustl.edu/). Next, the repetitions of the same voxel at given diffusion time and b-value range were averaged to obtain the mean diffusion signal decay that was used in further analysis.

#### 2.5.2 Estimation of the compartment size (*L*)

The free diffusion coefficient *D*_0_ and *L* were simultaneously estimated by fitting from Eqs. 1-2, 7 in the case of MC simulation (when longitudinal relaxation is neglected) and from Eqs. 2-3,7 in the case of the in-vivo and ex-vivo experiments. The fittings were conducted using Bayesian probability theory-based software.

### 2.5 Histopathologic assessment and slides analysis

Excised eWAT tissues were dehydrated in 70% ethanol before embedding in paraffin wax. Tissues were sectioned at a thickness of 5 µm and mounted on positively charged glass slides and then stained with hematoxylin and eosin (H&E). Brightfield whole-slide images were captured at 40x magnification using Hamamatsu NanoZoomer whole-slide imaging system. After whole-slide imaging, a region of 3.5 x 2 mm^2^ was analyzed per sample (number of cells>3000) using MIPAR Image Analysis Software version 3.0.3 (MIPAR Software LLC, Worthington, OH). The image analysis was as follows: 1. The single 2D black and white (BW) adipose tissue image was windowed and leveled manually to get the better image contrast. 2. A basic threshold (65%) was used to generate the label of adipocytes (white area inside the cell). 3. All the features’ area less than 50 pixel (<92 µm^2^) were removed. 4. To get better separation between the neighbor features the uniform shrinkage by 1pixel (1.85 µm) in all directions was performed. 5. The adipocytes with partially imaged volume on the edge of the 2D image frame were automatically removed. 6. The resulting cells maps of each subject were visually evaluated and manually corrected to eliminate any pixels outside the cells of interest and to include any cells that were not automatically segmented. All subjects were then processed using the recipe developed above on a single subject. At the end of analysis, the mean linear intercept lengths *L_m_* recorded from each cell was measured with number of lines set to 100. To estimate the *L*, we used the well-known relationship between the linear intercept length *L_m_* and the surface to volume ratio [38, 39]:

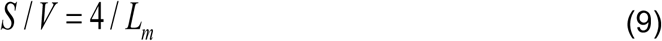

The size of spherical droplet is:

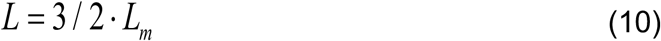

## 3. Results

To verify the validity of the “short-time-regime” method for determining the size of compartments filled with slowly diffusing particles, e.g., lipids, MC simulations were performed with varying SNRs (40, 60, 80, 100, and 200), and input *D* = 0.0175 µm^2^ / ms (characteristic of lipid diffusion at 37 °C), and *L* of 20, 30, 40, 50, 60, 80 and 100 μm (as mentioned above, no longitudinal relaxation is included). Representative simulated signal values at varying b-values and diffusion times follow the expected trend of a “shallower” signal decay with increasing diffusion times; an effect of longer diffusion times accommodating more chances for protons to encounter compartment wall and thus increasing the fractional population of “restricted” protons in the compartment (Fig. 1A). At SNR values of >40 simulated *L* s were reliably estimated but with an ∼8% underestimation (Fig. 1B). The underestimation of *D*_0_ at SNRs above 40 was estimated to be ∼2% (Fig. 1C).

**Figure 1.**
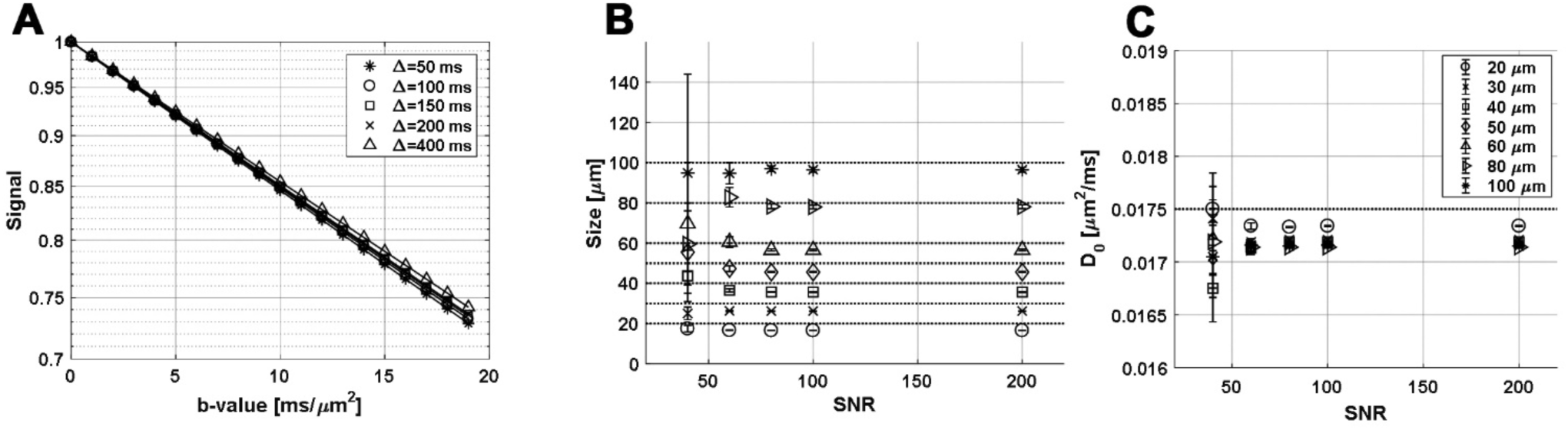
Simulation results (symbols) of diffusion-weighted MRI signal decay as a function of b-value at differing diffusion times, Δs, along with Bayesian probability Theory-based fits of Eq. 1 (solid lines) to the simulated data **(A)**. Data shown here are representative of a compartment with a size (*L*)of 50 μm. Estimated *L* s (± calculation uncertainties) calculated from Eqs. 1, 2 and 7 are compared to the simulation input *L* s for all SNR values greater than 40. In general, the estimated *L* is 8% smaller than the input (true) *L*. **(B)**. Estimated *D*_0_ was very close (<2.5% difference) to known *D*_0_ value of 0.0175 μm^2^/msec used for simulations for all SNR values greater than 40 **(C)**.

Representative simulated datasets in which calculated *L* is plotted as functions of true *L* at SNRs of 40, 60, and 80 are instructive (Fig. 2). *D*_0_ estimates are stable for all *L* sat SNRs greater than 40 (Fig. 2A, E, I). Here, we used a coefficient of 1.02 to correct for systematic 2% underestimation of *D*_0_ (Fig. 2B, F, J). The systematic 8% underestimation of *L* is made clear in plots of estimated versus input (true) *L*, as is the increase in parameter estimate uncertainty with a decrease in SNR (Fig. 2C, G, K). The systematic 8% underestimation of *L* is correctable with a coefficient of 1.08 (Fig. 2D, H, L).

**Figure 2.**
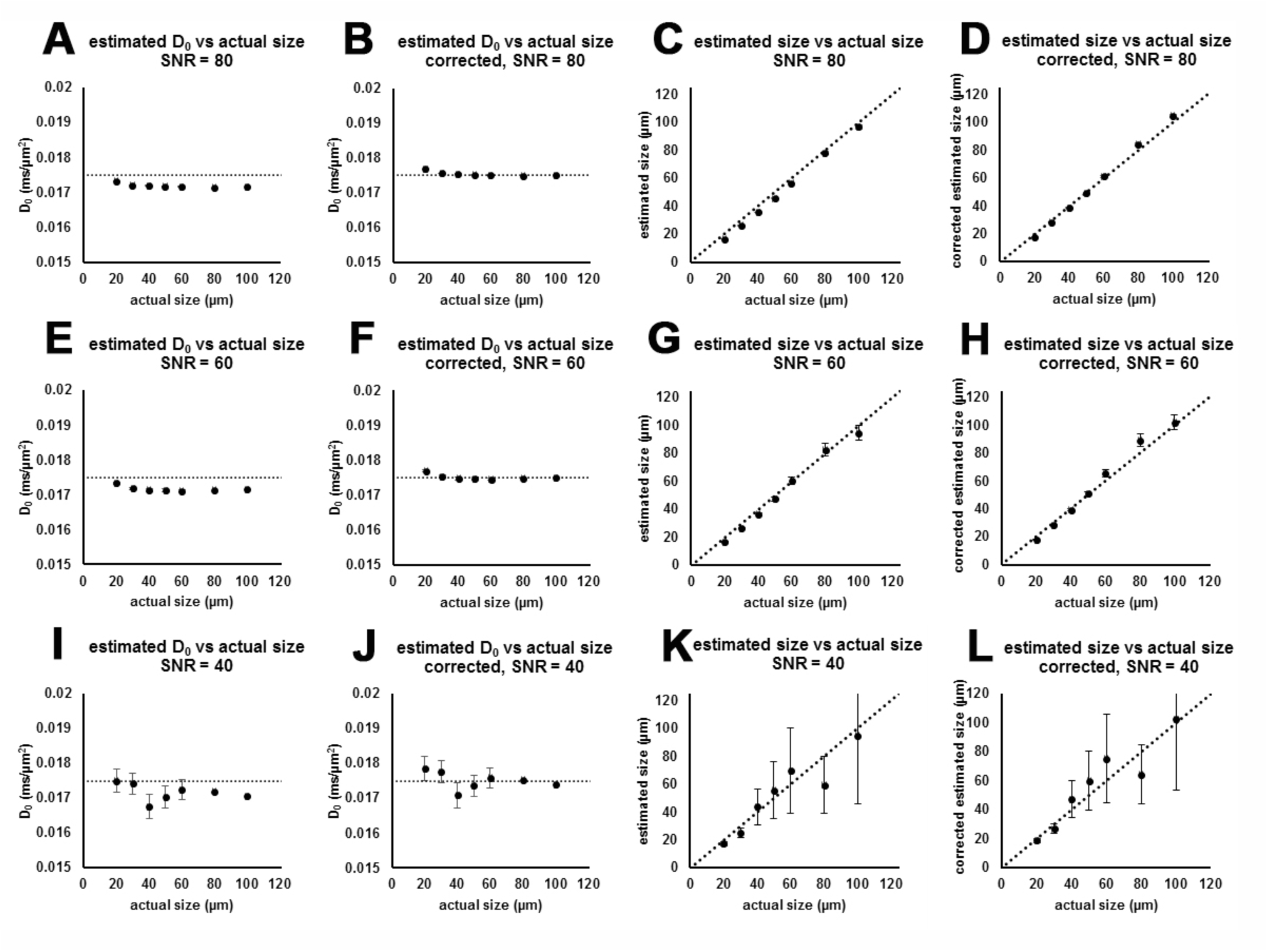
Calculated lipid diffusivity (**A, E, I**) and size (*L*) (**C, G, K**) at decreasing SNR values (top to bottom) estimated from Eqs. 1-2, 7 are compared against the true (input) *L* values. There is 2% and 8% underestimation of *D*_0_ and *L* using this method, respectively. Based on the systematic underestimation of *L*, a coefficient of 1.02 or 1.08 as an empirical correction to diffusivity or *L* estimation, respectively, can be used to make accurate diffusivity (**B, F, J**) and compartment *L* measurements (**D,H,L**).

### In-vivo and ex-vivo eWAT vs. histology

Diffusion-weighted spectroscopy data were collected at five different diffusion times between 50 to 400 ms and eight b-values, ranging from 1.5 to 20 ms/*μm*^2^ per diffusion time. A total of 40 diffusion-weighted datapoints were used to estimate adipocyte size. Data were collected both in vivo and ex vivo on the same fat pads (symbols, Figure 3C, D). *D*_0_, *R*_1_ and size (*L*) were simultaneously estimated by fitting using Eqs. 3 and 7 (dash lines, Figure 3C, D), adipocyte size was corrected using coefficient of 1.08 (see simulations above), and calculated MR-based adipocyte size were compared to adipocyte size distributions obtained from histological analysis (Figure 3F; Table 1).

**Figure 3.**
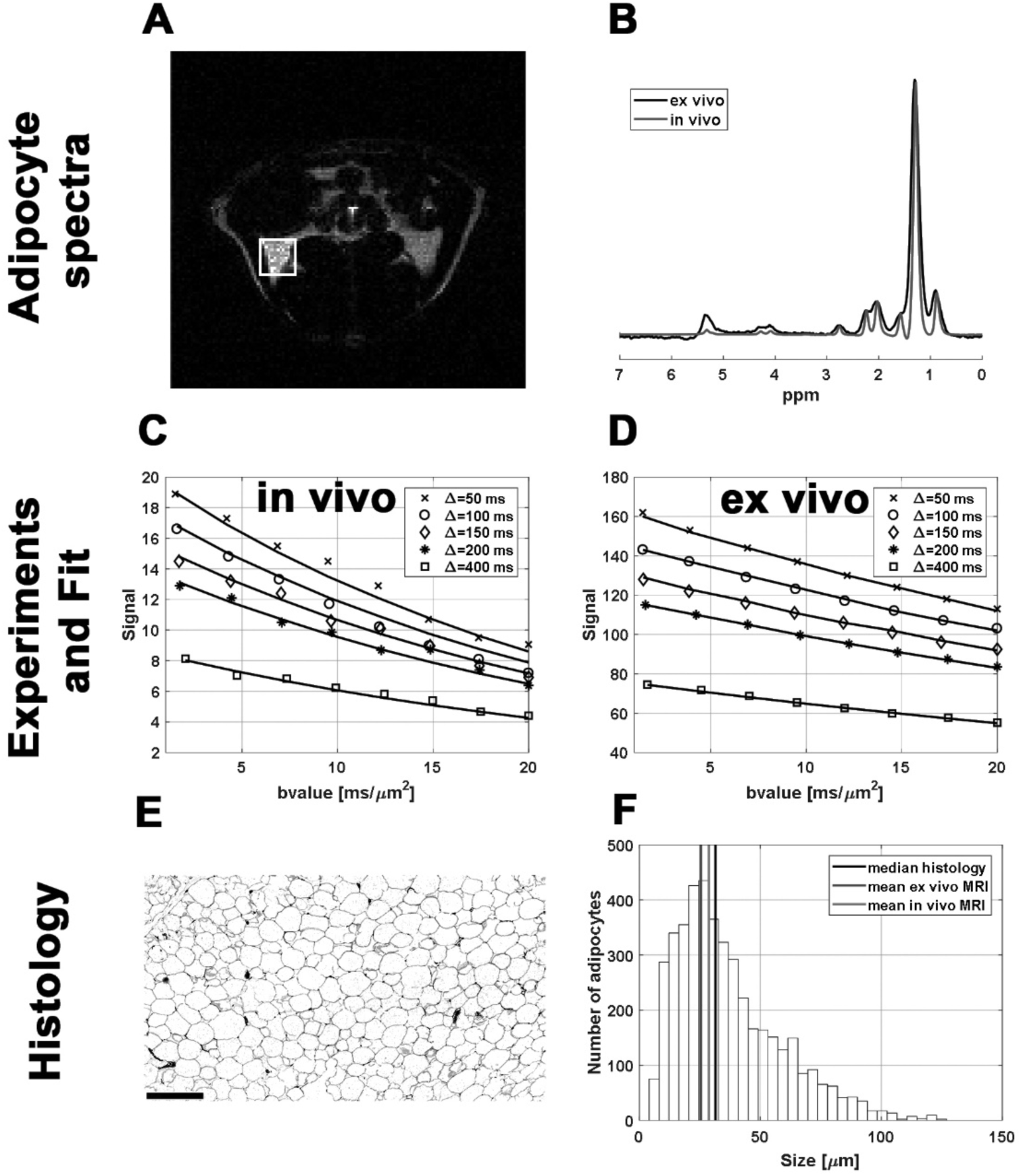
MR and histology results of Rat 1. A representative in vivo anatomical MR image of eWAT **(A)** and spectrum obtained from both in vivo (grey line) and ex vivo (black line) eWAT **(B)**. A water suppression pulse was used to emphasize fat regions within the rat body. The white box in (A) depicts the voxel region of the subsequent diffusion-weighted STEAM spectroscopy MR experiments. Signals decays vs. b-values obtained with diffusion times of 50, 100, 150, 200 and 400 ms and eight b-values (max b-value 20 ms/*μm*^2^) per diffusion time for a total of 40 diffusion-weighted datapoints for **(C)** in vivo and **(D)** ex vivo eWAT. Symbols represent experimental data and lines represent the fit of Eq. 3 to the data. The model parameters (the diffusion coefficient *D*_0_, droplet size *L*, longitudinal relaxation rate constant *R*_1_) were simultaneously estimated in the framework of the Bayesian probability analysis. Comparisons between droplet size distributions obtained from histology and size obtained from diffusion MRI is shown in (F). The black, gray and light gray vertical lines represent the median adipocyte size obtained from histology, and the mean size obtained from ex vivo and in vivo MRI, respectively.

**Table 1:** Adipocyte size (*L*) calculated from in vivo and ex vivo diffusion-weighted MR and histology.

| | MR-derived size ($\mu\text{m}$)<br>(calculated radius ± calculation uncertainty) | | | | Histology-derived size ($\mu\text{m}$)<br>(median; median abs deviation) |
| --- | --- | --- | --- | --- | --- |
|  | In vivo |  | Ex vivo |  |  |
| | size estimate ($\mu\text{m}$)<br>size uncertainty | Abs % deviation from histology | size estimate ($\mu\text{m}$) | Abs % deviation from histology | |
| Rat 1 | 28.7 | 1.4 | 25.4 | 12.7 | 29.1; 10.8 |
| Rat 2 | 25.7 | 9.2 | 32.3 | 14.1 | 28.3; 13.2 |
| Rat 3 | 25.1 | 38.0 | 34.7 | 14.3 | 40.5; 16.1 |
| Mean ± StDev | 26.5 ± 1.6 | 16.2 ± 15.8 | 30.8 ± 3.9 | 13.7 ± 0.7 | 32.6 ± 5.6 |

## 4. Discussion

In this study we developed a direct, non-invasive method to measure adipocyte size using short-diffusion-time-regime MR measurements. The accuracy and limitation of our approach for cell size estimation was first examined using Monte Carlo simulations of “randomly-walking” particles within spheres of different sizes. The method was then applied to estimate the sizes of epididymal white adipocytes in vivo and ex vivo and compared against gold-standard histology measurements.

To determine how compartment size estimation depends on *b*-value, diffusion time, and SNR, Monte-Carlo simulations were performed with *D*_0_ similar to that of lipids at 37°C. For accurate estimation of size two following conditions should be met. The first condition is the validity of short-time approximation that requires the second term in Eq. 4 to be much smaller than 1. In other words, the first condition that should be met is:

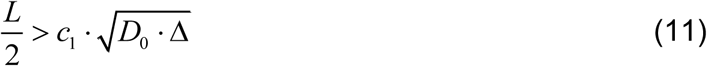

Moreover, to distinguish *D*_0_ and *D*(*t*), the difference between correspponding signals should be bigger than the noise level, leading to second condition that should be met:

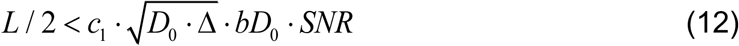

Our simulations and aforementioned conditions revealed that, in order to make an accurate estimation of sizes larger than 50 µm, *SNR* > 100 should be achieved (data not shown). This condition can be ensured with high quality transmit/receive hardware, larger voxel sizes, and relatively short echo times. Note also that there is the systematic underestimation of size, which can be corrected by applying the empirical coefficient 1.08 (Fig. 2C, F). This coefficient is universal and its underlying biophysical basis will be the subject of future investigation.

In vivo and ex vivo MR-based estimates of adipocyte size appear to be in the reasonable agreement with the histology-based measures. It was demonstrated that even after applying the empirical coefficient 1.08, a slight bias towards underestimating adipocyte size remains both in vivo and, more pronounced, in in vivo measurements. We hypothesize that this underestimation is caused by the substantial difference in SNR between in vivo and to ex vivo acquisitions: *SNR* ∼ 150 for in vivo and *SNR* ∼ 2500 for ex vivo. It also might be originated from slight epididymal fat motion that may occur due to pulsation of surrounding organs. This limitation might be partially resolved in clinical MRI studies of epididymal fat, where the diffusion experiments can be conducted with breath hold and cardiac gating.

In brief, the short-time-regime model relating adipocyte size and the diffusion signal can be effectively used for an accurate estimation of compartment size. It should be noted, however, that the estimation becomes less accurate when the droplet size increases because the deviation of *D*(*t*) from *D*_0_ becomes smaller (it is inversely proportional to *R*, see the second term in Eq. 7), when the fractional population of lipids interacting with a droplet boundary *decreases* as the droplet size *increases*. This problem is especially pronounced when SNR decreases. Moreover, for sufficiently low SNR, the difference can be completely “lost” due to the noise.

The short diffusion time MRI approach presented in this study can be used as a “non-invasive histology tool” to study health and disease of abdominal fat. The proposed diffusion MRI approach is relatively rapid and doesn’t require complex biophysical modeling of the data. In addition, due to the use of relatively low diffusion weighting the proposed approach might have clinical potential. While this work has focused on measuring the sizes of large lipid droplets found in white adipocytes (which allowed us to estimate adipocyte size), our method may be even better suited for measurement of smaller lipid droplets found in brown adipose tissue or fatty liver which can range in size from less than 1 µm to tens of µm. For example, this MR-based measure of adipocyte size would be a powerful tool in monitoring the utilization/consumption of lipid during brown adipose tissue activation via the shrinkage of lipid droplet in the brown adipocytes [40–44]. Similarly, this method could be used to longitudinally study and diagnose microvascular steatosis in the liver, a condition which is correlated with severity of non-alcoholic fatty liver disease.

It is important to recognize that variance in temperature, lipid viscosity, and lipid chemical composition will affect the free diffusivity of lipids and thus propagate into the measures made herein if not accounted for. To resolve this issue, we opted to treat the free diffusivity of lipid, *D*_0_, as unknown and calculate its value simultaneously with size (Eq. 7). For this purpose, diffusion-weighted MR data were collected using multiple *b*-values *and* diffusion times Δ, for a total of 40 datapoints over two dimensions (eight b-values for each of five different Δ).

It should also be mentioned that we here report only a proof-of-concept and that further study is needed to optimize the proposed method and explore its limits. For example, the next logical step would be to apply it to mouse models of obesity and compare against healthy lean mice. Such a study would provide a real-world description of the methods ability to discern health from disease. Another option is to use human subcutaneous biopsies from lean and obese subjects to evaluate the robustness of the method to serve as a “non-invasive biopsy” approach to study adipose tissue heath.

Our model may (and should) be enhanced/optimized. Firstly, it will be important to optimize the number of b-values and diffusion times acquired to provide robust estimates of size. For example, the number of diffusion times should likely be increases (at the expense of the number of b-values) for the purpose of optimizing the simultaneous estimation of diffusivity and size. The model may also be improved by incorporating statistical distributions of adipocyte size revealed in the histological data (e.g., the gamma-distribution).

## Conclusion

Diffusion MR is shown to be a potential tool for direct, non-invasive, and quantitative measurement of adipocyte size. Our approach may be useful for longitudinal studies of type 2 diabetes, brown adipose tissue function, steatohepatitis, and fatty liver disease.

